# Learning-induced sequence reactivation during sharp-wave ripples: a computational study

**DOI:** 10.1101/207894

**Authors:** Paola Malerba, Katya Tsimring, Maxim Bazhenov

## Abstract

During sleep, memories formed during the day are consolidated in a dialogue between cortex and hippocampus. The reactivation of specific neural activity patterns – replay – during sleep has been observed in both structures and is hypothesized to represent a neuronal substrate of consolidation. In the hippocampus, replay happens during sharp wave – ripples (SWR), short bouts of excitatory activity in area CA3 which induce high frequency oscillations in area CA1. In particular, recordings of hippocampal cells which spike at a specific location (‘place cells’) show that recently learned trajectories are reactivated during SWR in the following sleep SWR. Despite the importance of sleep replay, its underlying neural mechanisms are still poorly understood.

We developed a model of SWR activity, to study the effects of learning-induced synaptic changes on spontaneous sequence reactivation during SWR. The model implemented a paradigm including three epochs: Pre-sleep, learning and Post-sleep activity. We first tested the effects of learning on the hippocampal network activity through changes in a minimal number of synapses connecting selected pyramidal cells. We then introduced an explicit trajectory-learning task to the model, to obtain behavior-induced synaptic changes. The model revealed that the recently learned trajectory reactivates during sleep more often than other trajectories in the training field. The study predicts that the gain of reactivation rate during sleep following vs sleep preceding learning for a trained sequence of pyramidal cells depends on Pre-sleep activation of the same sequence, and on the amount of trajectory repetitions included in the training phase.

## Introduction

Memories are composed of three stages: acquisition, consolidation and retrieval. The consolidation phase provides memory resilience to interference, and is influenced by sleep. Indeed, memory performance benefits from sleep (Mednick, Nakayama et al. 2003, Mednick 2009, Mednick, Cai et al. 2011, Wilhelm, Diekelmann et al. 2011, Rasch and Born 2013), and specific sleep features have been linked to increased memory performance (Marshall, Helgadottir et al. 2006, Mednick, McDevitt et al. 2013, Rasch and Born 2013, McDevitt, Duggan et al. 2014). Sleep is a stage in which the brain can be described as self-organizing, because it is not processing external inputs. During sleep, the local average activity of brain regions can be measured to find characteristic oscillations which vary across frequency, brain regions and time scales. Furthermore, experimental results show that the coordination of brain rhythms during sleep promotes (and possibly mediates) memory consolidation both in humans and animals (Staresina, Bergmann et al. 2015, Latchoumane, Ngo et al. 2017). In particular, sharp-wave ripples (SWR) are rhythms present in wake and sleep in the hippocampus, a brain region which is crucial in encoding and retrieving recently formed memories (Buzsaki, Buhl et al. 2003, Schwindel and McNaughton 2011, Buzsaki 2015). A SWR complex is formed by the combination of two events, both occurring simultaneously in different layers of the Local Field Potential (LFP) of hippocampal area CA1. 1) In stratum radiatum, a large deflection (the sharp wave) lasting 100-200 ms is caused by a barrage of excitatory inputs coming from area CA3, where a large spiking event among the highly interconnected pyramidal cells generates intense drive for area CA1 cells (both pyramidal and interneurons). 2) In stratum pyramidale, high-frequency (>150 Hz) short-lived (50-80 ms) oscillations (the ripple) are seen as a result of the CA3 input. In fact, the local fast-spiking interneurons of CA1 are known to fire at high frequency for the duration of a ripple, so their spikes are hypothesized to pace the oscillation. During a ripple, CA1 pyramidal cells are receiving excitation from CA3 and inhibition at ripple frequency from CA1 interneurons; as a result most of them are suppressed, and the few (about 10% (Csicsvari, Hirase et al. 1999)) that spike do so within windows of opportunity left by the local interneurons.

During learning, some CA1 pyramidal cells that fire in a specific location as the animal explores an environment have been labeled ‘place cells’ (O'Keefe 1976, O'Keefe and Nadel 1978, O'Neill, Senior et al. 2006); and it has been shown that cells which are active together during learning (e.g. place cells that code for nearby locations explored during a recently learned task) also activate in the same SWR during sleep (Wilson and McNaughton 1994, Skaggs and McNaughton 1996, Sutherland and McNaughton 2000). Furthermore, if the task involves learning a specific path and the spikes of place cells along that path are recorded, the sequence of spikes among those cells is reactivated in the correct order (in a time-compressed manner) during SWR in the subsequent sleep epoch (Kudrimoti, Barnes et al. 1999, Girardeau and Zugaro 2011). A specific path learned during a task can then be seen reactivated (in a time-compressed manner) during both awake and sleep SWRs (Nádasdy, Hirase et al. 1999). Sequence reactivation during SWR in sleep directly affects memory: both the spike sequences within SWR and the number of SWR during sleep correlate with memory performance, and suppression of sleep SWR impairs memory consolidation (Girardeau, Benchenane et al. 2009, Ego-Stengel and Wilson 2010). Hence, it is crucial to explain how spiking activity related to learning enables the reactivation of the correct spike sequences in SWR during sleep.

In general, the influence of learning over sleep reactivation content is performed by comparing spiking in the nights before (Pre-sleep) and after (Post-sleep) a learning paradigm. When place cells which code for locations that are relevant to the learned task reactivate more strongly in Post-sleep compared to Presleep, it is safe to state that learning the task has influenced the spiking content of SWR reactivation. The specific mechanisms mediating such influences are yet to be determined. Common hypotheses involve synaptic plasticity and neuromodulators, and the overall theory states that during learning (and possibly during awake SWRs) synapses in the hippocampus are changed, and those changes induce the enhanced representation of the learned spiking sequences in SWR during Post-sleep.

*In vivo* studies have shown that during sleep, pyramidal cells can be divided in those involved in many SWR and those mostly not spiking in SWR, and this separation seems to persist across days (Csicsvari, Hirase et al. 1999, Csicsvari, Hirase et al. 2000, Buzsaki 2015). Furthermore, in a Presleep/experience/Post-sleep paradigm, Grosmark et al. (Grosmark and Buzsaki 2016) have shown that pyramidal cells can be separated into “rigid” and “plastic”. The rigid group has a higher firing rate, low spatial specificity and shows very little change across the sleep/experience/sleep paradigm. The plastic group has a lower firing rate, but shows increasing spatial specificity and increasing ripple reactivation during the experience phase of the paradigm. The plastic cells are those who go on to show increased bursting and co-activation during SWR in the Post-sleep phase of the paradigm. In this work, we investigated how synaptic plasticity can influence spontaneous reactivation of spike sequences during sleep in our previously developed biophysical model of SWR activity in a Pre-sleep/learning/Post-sleep paradigm. The effect of learning on cell reactivation was first studied in a simplified representation, where we manually changed very few synapses, and then in a learning paradigm that was explicitly modeled to represent a “virtual rat” exploring an environment, within which a specific trajectory was rewarded (and hence learned). The spike times obtained for the training phase were used to induce offline spike-timing dependent plasticity (STDP) in synapses among cells in the network, leading to different spiking profiles in Pre-sleep and Post-sleep simulations. We find that learning-dependent plasticity is able to enhance the representation of cell sequences (and hence space trajectories) during spontaneous SWR. Our model predicts that this enhancement depends on the co-activation in the Pre-sleep epoch, the timing (within SWR events) of the first spike of the spike sequence and the amount of training included in the learning phase of the experiment.

## Results

### Changes in minimal number of synapses can promote reactivation of cell triplet in CA3

In this study, we started from our previously developed computational model of stochastic, randomly emerging SWR (Malerba, Fodder et al. 2016, Malerba, Krishnan et al. 2016). This model includes two hippocampal regions, CA3 and CA1 (Fig 1A), each represented with pyramidal cells (excitatory) and basket cells (inhibitory interneurons). Area CA3 has highly recurrent excitatory connectivity (Shepherd 2004) and sends excitatory projections to area CA1 (the Schaffer Collaterals), reaching both excitatory and inhibitory neurons (Shepherd 2004). Within CA1, the excitatory connections between pyramidal cells are very few and sparse, as shown by experimental results (Deuchars and Thomson 1996). Parameters for cells in our model are chosen so that: 1) pyramidal cells have physiological firing rates and show bursting (Andersen, Morris et al. 2006), and 2) basket cells are fast-spiking neurons, with very low spike-frequency adaptation, and are then capable of spiking at high frequencies in response to sustained strong inputs (Andersen, Morris et al. 2006). While the rules shaping network connectivity stayed the same, multiple instances of specific connectivity matrices were used, to represent different “virtual rats” in our computational study.

**Figure 1.**
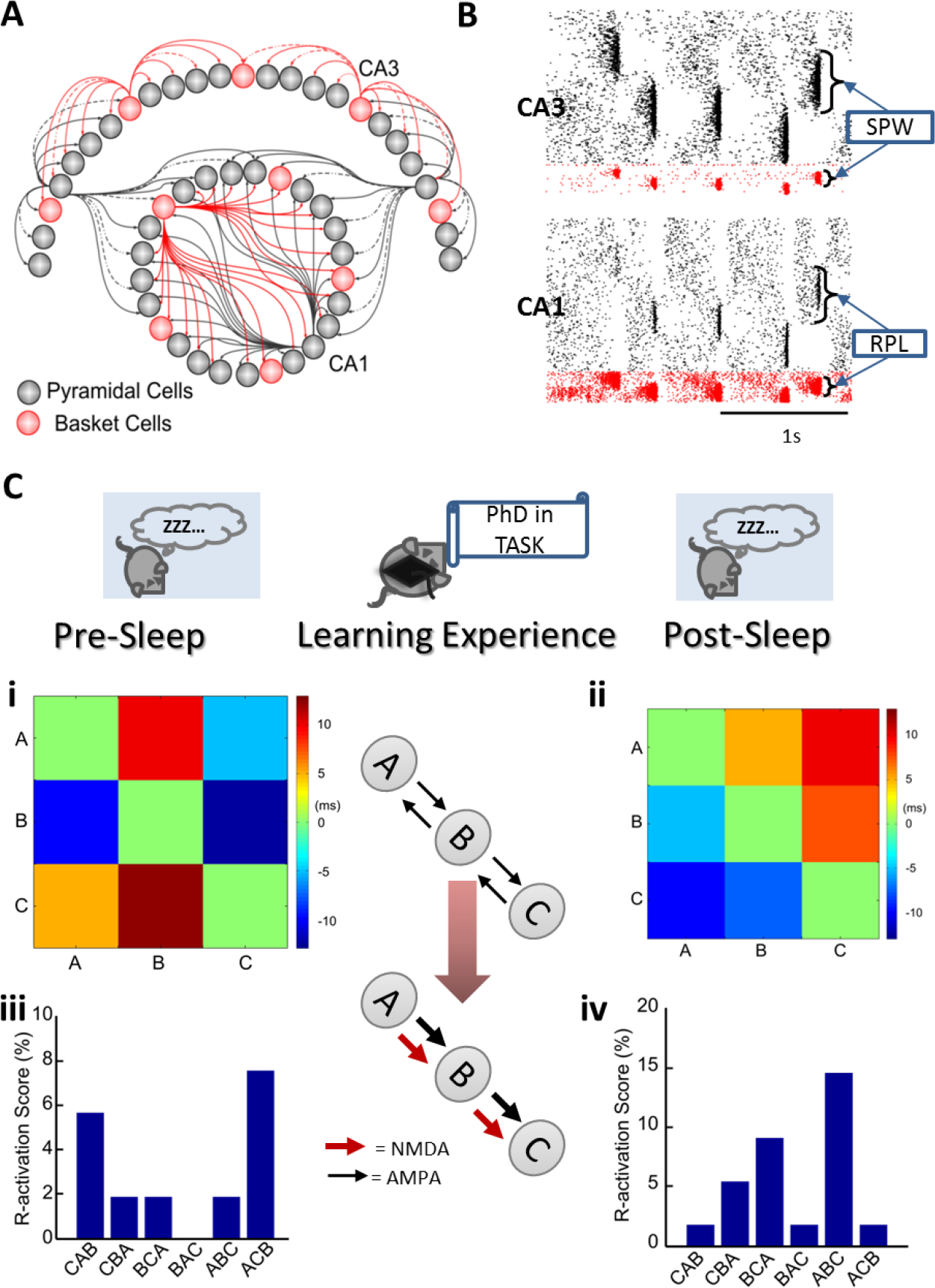
Changes in minimal number of synapses can promote reactivation of cell triplet. **A.** Model schematic representation. The model includes 1200 pyramidal cells and 240 interneurons in CA3, and 800 pyramidal cells and 160 interneurons in CA1. Connections within CA3 include high recurrence among pyramidal cells, and a topological preference for neighbors within a radius of about one third of the network. CA3 pyramidal cells project to CA1 cells (both pyramidal cells and interneurons). Within CA1, the all-to-all connectivity is not effective between CA1 pyramidal cells, which are weakly and sparsely connected. **B.** Example of model network SWR activity. Noise-driven spiking in CA3 occasionally triggers an excitatory cascade of spikes in pyramidal cells and interneurons (the sharp wave,SPW). Spiking of CA3 pyramidal cells drives CA1 interneurons to spike in short-lived high-frequency ripples (RPL), while CA1 pyramidal cells receive competing excitation from CA3 and later inhibition from CA1 interneurons. a few CA1 pyramidal cells are driven to spike within windows of opportunity left by the rhythmic local inhibition. **C.** Representation of the main question addressed in this work: Comparing the reactivation of cell sequences in two sleep epochs, and study the role of learning in shaping the difference. In our model, we represent learning by altering synaptic connections as shown in the middle plot: AMPA synapses promoting the spiking order “ABC” are increased, and the AMPA synapses promoting the opposite spike order are removed. In addition, NMDA synapses in the direction favoring “ABC” spiking are introduced. Data from one example simulation is used to show how spiking changes in Post vs Pre sleep simulations. Matrices shows an example of spike time differences between three selected pyramidal cells in the CA3 network in a Pre-sleep simulation (Ci) and a Post-sleep simulation (Cii). The bar plots (Ciii-iv) show the R-activation score (% of ripples in which a given triplet spiked) for the ordered triplet “ABC” and all its permutations. Note that the “ABC” order R-activation is larger for Post-sleep than it is for Pre-sleep.

In our model, every cell was in a noise-driven spiking regime (as opposed to a limit cycle periodic spiking regime), a feature which introduced random fluctuations in the background network activity. Recurrence within excitatory neurons in CA3 was the gateway for occasional large excitatory spiking events (the sharp waves, SPW) which were projected to CA1 interneurons and pyramidal cells (Fig 1B). Concurrently, the drive from CA3 sharp wave spiking imposed high-frequency firing on the basket cells of area CA1, which formed the structure for the ripple, and determined the ripple frequency. All the while, CA1 pyramidal cells received excitation from CA3 and inhibition from CA1 basket cells. Since CA1 pyramidal cells lack recurrent excitability to organize their firing, the competition between these inhibitory and excitatory inputs only allowed a small percentage of pyramidal cells in CA1 to spike during a ripple, with timing controlled by windows of opportunity left by the ongoing inhibitory ripple oscillations (Fig 1B). Note that in our model, sharp waves in CA3 did not invade the totality of the network, but were instead localized. This is consistent with *in vivo* recordings showing that different simultaneous channels capture SWR activity differently (Patel, Schomburg et al. 2013). Localized sharp wave activity in CA3 drove localized ripple activity in CA1, as can be seen in Fig 1B. In this work, we took advantage of the biophysical model of SWR spontaneous activity during sleep to analyze the role of changing excitatory synapses in shaping the activation of cell sequences across SWRs. This study centered on CA3 cells and their spiking activity during SWRs.

The main setup of this study (Fig 1C) was to compare two model simulations, one representing Pre-sleep (the sleep epoch preceding training) and one representing Post-sleep (the sleep event following training). If the specific network connectivity, parameters and input-noise traces were kept the same in the Pre and Post-sleep simulations, the two sleep events would look identical. We imposed offline synaptic modifications between Pre and Post-sleep simulations to represent the effects of a learning experience and then run a Post-sleep simulation where every input and connections were copied from the Pre-sleep simulation, except for the learning-modified synapses. To quantify how often any cell sequence reactivated spontaneously across SWR in a simulation, we defined its “Ripple-activation score” (R-activation score) as the percentage of SWR during which all cells in the sequence spiked in the correct order. In a first simplified setting, we chose three CA3 pyramidal cells (cells A, B and C) to form an ordered triplet (our most simple cell sequence) and found their R-activation score during the Pre-sleep simulation (in Fig 1C iii, one example of ABC R-activation score is shown together with the R-activation scores of all its permutations). We then altered the excitatory synaptic connections between the ABC cells as follows (Fig 1C, center panel): we found the strengths of AMPA (short-lived) excitatory synapses which connect A to B and B to C, and replaced their synaptic strengths with the maximum value across all the CA3-CA3 pyramidal cell synapses in the network. We also introduced NMDA (long-lasting) excitatory synapses between A to B and B to C cells. Finally, we removed (if present) the excitatory AMPA synapses from B to A and from C to B. It is to note that in Pre-sleep there are no NMDA synapses in the network, so we did not need to remove any of them. In the example of Fig 1C, this artificial change in very few synapses in the network led to a strong increase in the R-activation score of ABC in the Postsleep simulation (Fig 1C iv). The ability of the new synapses to improve the reactivation of the ABC spiking sequence was further supported by the comparison between the average spike time differences between cells of the triplet in Pre-sleep (Fig 1C i) and Post-sleep (Fig 1 C ii).

### Pre-sleep activation modulates learning-induced gain in Post-sleep reactivation

Using the setup we developed in the previous section, we could represent activity in the Pre and Postsleep phases and quantify the change in sequence reactivation between the two sleep stages due to synaptic changes introduced during learning. Across multiple Pre-sleep simulations (n=17), we found an inherent range of R-activation scores for ordered cell triplets (Fig 2A) which, surprisingly, was not uniformly distributed among triplets. In fact we could fit a log-normal curve to the distribution, in agreement with numerous experimental observations (Buzsaki, Buhl et al. 2003, Buzsaki 2015), which also emphasizes that this type of variance in cells firing rates and R-activations could optimize coding strategies for the hippocampal system. We chose to study the effects of changing few synapses over a pool of ABC triplets spanning 0% to 25% R-activation scores in the Pre-sleep simulation to cover most of the range of R-activations found in the distribution (see Computational Methods for details on how the cells composing each triplet were chosen). Each simulation/triplet pair (which can be thought of as representing a subject/task pair in an experimental study) then underwent the synaptic modifications introduced in Fig 1C: maximized AMPA synapses favoring the triplet order, removing AMPA synapses opposing the triplet order and introducing NMDA synapses along the maximized AMPA ones. For each simulation/triplet case we then compared Pre and Post-sleep R-activations of the ABC triplet. We call the difference between the Post-sleep score and the Pre-sleep score for a triplet its “Score Gain”. In Fig. 2B, we show that the Score Gain of a triplet correlated with its Pre-sleep R-activation, and hence when this simplified learning was applied to a triplet with too high R-activation during Pre-sleep it could not increase its R-activation, or might even have decreased it. This is intuitively coherent with some kind of ‘ceiling effect’, in which Post-sleep R-activation could only reach a preset amount (possibly limited by the overall spontaneous network activity, which is imposed by average network properties) and hence a Pre-sleep R-activation too high limited the available range for Gain.

**Figure 2.**
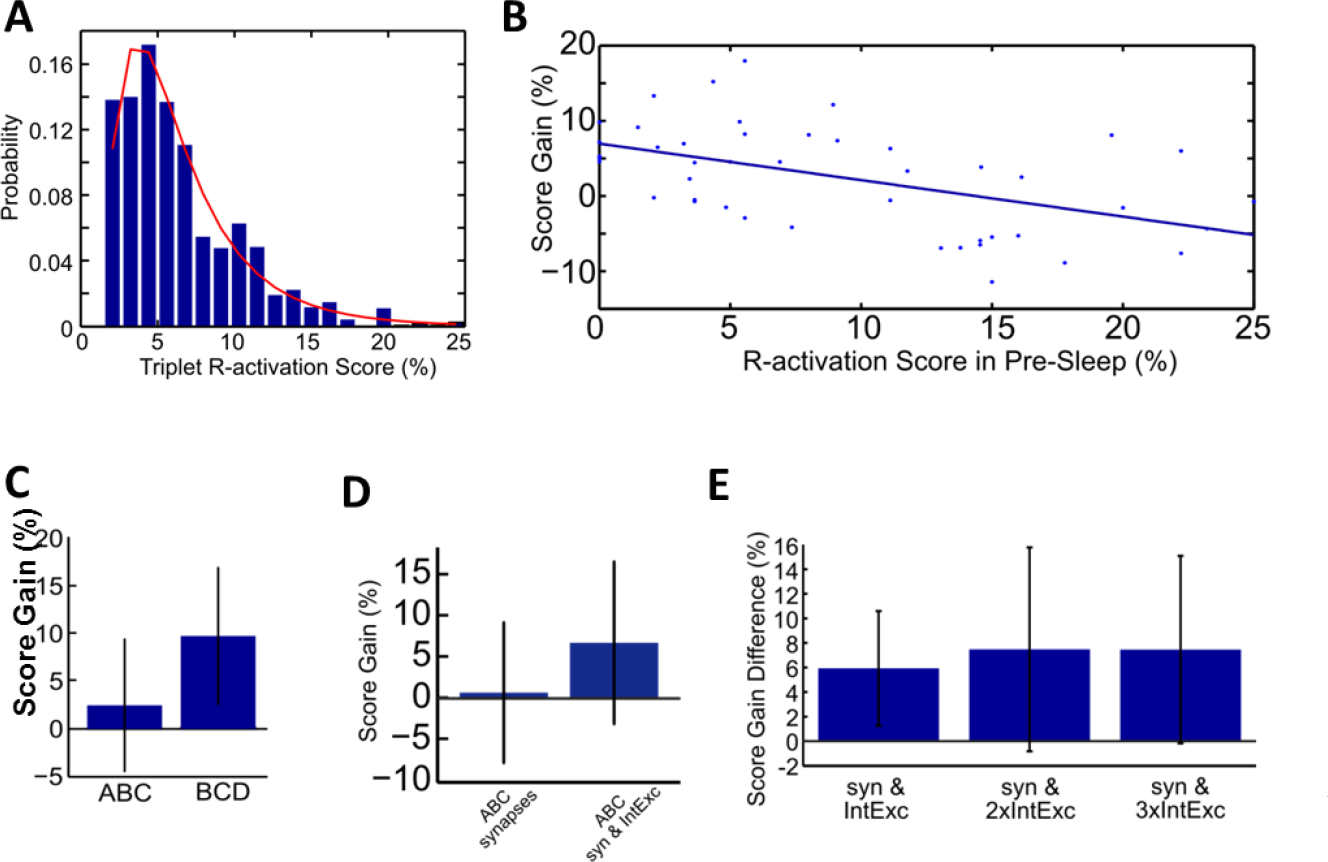
Pre-sleep activation modulates learning-induced gain in Post-sleep reactivation. **A.** Distribution of triplet reactivation scores across 17 simulations lasting 50s. The red line shows a log-normal distribution fit to the data (mean=1.6921 with 95% CI 1.6621 to 1.722, standard deviation=0.6125 with 95% CI 0.5921 to 0.6345). **B.** Gain in post-sleep reactivation is higher for triplets with lower pre-sleep activation. Each dot is a separate simulation pair, in which a triplet “ABC” R-activation is found in the Pre-sleep (x-axis value) and in the Post-sleep. The y-axis shows the difference between Post-sleep and Pre-sleep R-activation score. The line shows a linear fit which emphasizes the statistically significant negative correlation (r=-0.5101, p = 3.443e-4)). **C.** Gain in Post-sleep R-activation increases if the first cell of the triplet receives extra synaptic input. Comparing triplet “ABC” and “BCD” in simulations where the changes in synapses was applied along the word “ABCD”. Hence triplet “BCD” has all cells receiving extra synaptic input, while in triplet “ABC” only the last two cells do. Plot shows the average Post-Pre R-activation score gain across 15 simulation tests, lasting 50s, and error-bars mark standard deviations. Having synaptic input on the first cell of the sequence gives an extra gain to the R-activation in Post-sleep (paired Student’s t-test p=1.9071 e −06). **D.** Gain on Post-sleep reactivation can increase with increased intrinsic excitability of first cell in triplet. The bar plot shows the gain in R-activation score (Post-sleep minus Pre-sleep) for triplet “ABC” when only synapses are manipulated (left bar) compared with the case in which synapses and intrinsic excitability of cell “A” are manipulated to represent learning. Error bars mark standard deviation. Note that on average the gain is higher for triplets with enhanced intrinsic excitability in the first cell (paired student’s t-test p=0.003). In this case we could only use triplets in which the first cell did not have a high intrinsic excitability in Pre-sleep (artificially increasing intrinsic excitability can lead to the cell spiking behavior changing from noise-driven to DC-current driven, and hence the cell spikes continuously and decoupled from network activity) (10 simulations used). **E.** If the increase in intrinsic excitability is too large, no further gain is introduced. Bar plot shows the difference introduced by additional intrinsic excitability to cell “A” compared to Post-sleep with only synaptic manipulation. Error bars mark standard deviations. Each triplet received a fix (equal) amount of increased excitability and such amount could be doubled (2xInt.Exc) or tripled (3xInt.Exc). Note that while the first bar shows that additional excitability to cell “A” can lead to increased gain (as compared to a case where only synapses are manipulated) the range of efficacy of such increase is small. In fact, additional increase in excitability does not reflect in additional gain in Post-sleep R-activation. Statistically, the first bar is different from zero (p=0.003), while the other two are not (2xInt.Exc p=0.0191, 3xInt.Exc p=0.0129, Student’s t-test)

In this simplified representation of a learning effect on synapses, it is important to note that the first cell in the triplet (cell “A”) does not receive any change in its input in Post-sleep compared to Pre-sleep. In some sense, this procedure could be overlooking the very beginning of the sequence to be reactivated, and that could be the reason why there was such a hard ceiling effect on the R-activation Gain for triplets in Fig 2B. To address this limitation, we first “prolonged” our test set, introducing a fourth cell to the sequence undergoing artificial synaptic manipulation. The rules for changing synapses stayed the same: maximize AMPA connections in the direction of the ordered sequence (A to B, B to C and C to D), remove AMPA connections opposing the ordered sequence (B to A, C to B and D to C) and introduce NMDA connections where AMPA connections were maximized. In this case we could compare the R-activation Score Gain of two triplets within each test: ABC and BCD. The first triplet, just like in Fig 2B, would not receive any enhancement to its first cell, but for the second triplet (BCD), its first cell would be receiving enhanced excitatory synaptic input, both fast (AMPA) and slow (NMDA). In Fig 2C we show that extra excitatory synaptic input to the first cell did provide an advantage in Score Gain by comparing the Score Gains of the two different triplet types across 15 simulations. It is to note that the two distributions were significantly different (paired Student's t-test p=1.9e-6), which implies that input to the first cell in the triplet is capable of driving Post-sleep enhanced activation. In line with this observation, we next tested whether enhancing the changes of R-activation of the first cell in the triplet by means other than synaptic excitatory input could still introduce a favoring bias in Score Gain for a triplet. In fact, it is known that cells of the same type in the same region still show a range of firing rates (often log-normally distributed (Mizuseki and Buzsaki 2013)). This heterogeneity was introduced in our model via a direct current parameter representing a cell’s “intrinsic excitability” (*I_DC_* in the equation for membrane voltage, see Computational Methods). It is known that different mechanisms can possibly alter a cell baseline (i.e. in the absence of additional input) firing rate and behavior, such as as effects of the neuromodulators on conductances (Vassalle 1987, McCormick, Pape et al. 1991, McCormick 1992) or the balance of ionic concentrations (Offner 1991). To test if a learning-induced change in intrinsic excitability of the first cell in the triplet could compensate for the lack of additional synaptic input, we compared 10 simulations in which the same triplet underwent only synaptic changes (same paradigm as for Fig 2B) or received a small increase of the intrinsic excitability in addition to the synaptic changes. In Fig 2D, we show that a small increase in synaptic excitability of the first cell in the sequence led to enhanced Score Gain from Pre-sleep to Post-sleep R-activation of a triplet. When we increased the intrinsic excitability amount added to cell “A”, we found (Fig 2E) that the effect was quickly saturated such as doubling or even tripling the increase could not significantly increase the Score Gain from Pre-sleep to Post-sleep. This suggests that while increased excitability in the first cell could contribute to enhancing Score Gain, the overall Post-sleep R-activation was not fully dominated by any one given factor, such as synaptic excitatory inputs or intrinsic excitability, but rather established by their interaction with the overall network activity (and hence the Pre-sleep R-activation).

### Simplified learning extends word length reactivation

Within the same simplified approach of representing the effect of learning over hippocampal network connectivity by manually changing a small set of hand-selected synapses, we extended our analysis to sequences of spikes longer than a triplet. We chose a length of seven cell-spikes to build our sequence, as SWR are short-lived events and fitting longer sequences within a single event would be hard. In fact, there is an ongoing hypothesis that memories of paths which would require a long sequence of many place cells are reactivated across multiple SWR which happen in a very quick sequence (SWR packets (Buzsaki 2015)). While our computational model shows SWRs happening at times in groups interspersed by long pauses (data not shown), in this work we focus on characterizing how changes in synapses affect changes in R-activation across many SWR, and longer sequences would introduce an ulterior complexity, possibly masking relevant effects.

We represent a sequence of 7 cells with the 7-letter ‘word’ ABCDEFG. In Fig 3A we show one example of the changes introduced in synaptic connectivity by our artificial learning representation. In comparing the matrices of synaptic connection weights between the cells representing ABCDEFG in the network, we maximized the connections favoring the direction of ordered reactivation, which resulted in a high synaptic strength in the upper diagonal of the Post-sleep synaptic connections matrix in Fig 3A, and removed the connections favoring the opposite reactivation order, which resulted in all zeros in the lower diagonal of the Post-sleep synaptic connections. Note that all other connections between cells in the sequence were left unchanged. Furthermore, the middle schematic in Fig 3A shows that NMDA connections were also attributed to the synapses which had maximized AMPA connection weights (further details on the composition of the sequence of 7 cells are reported in Computational Methods).

**Figure 3.**
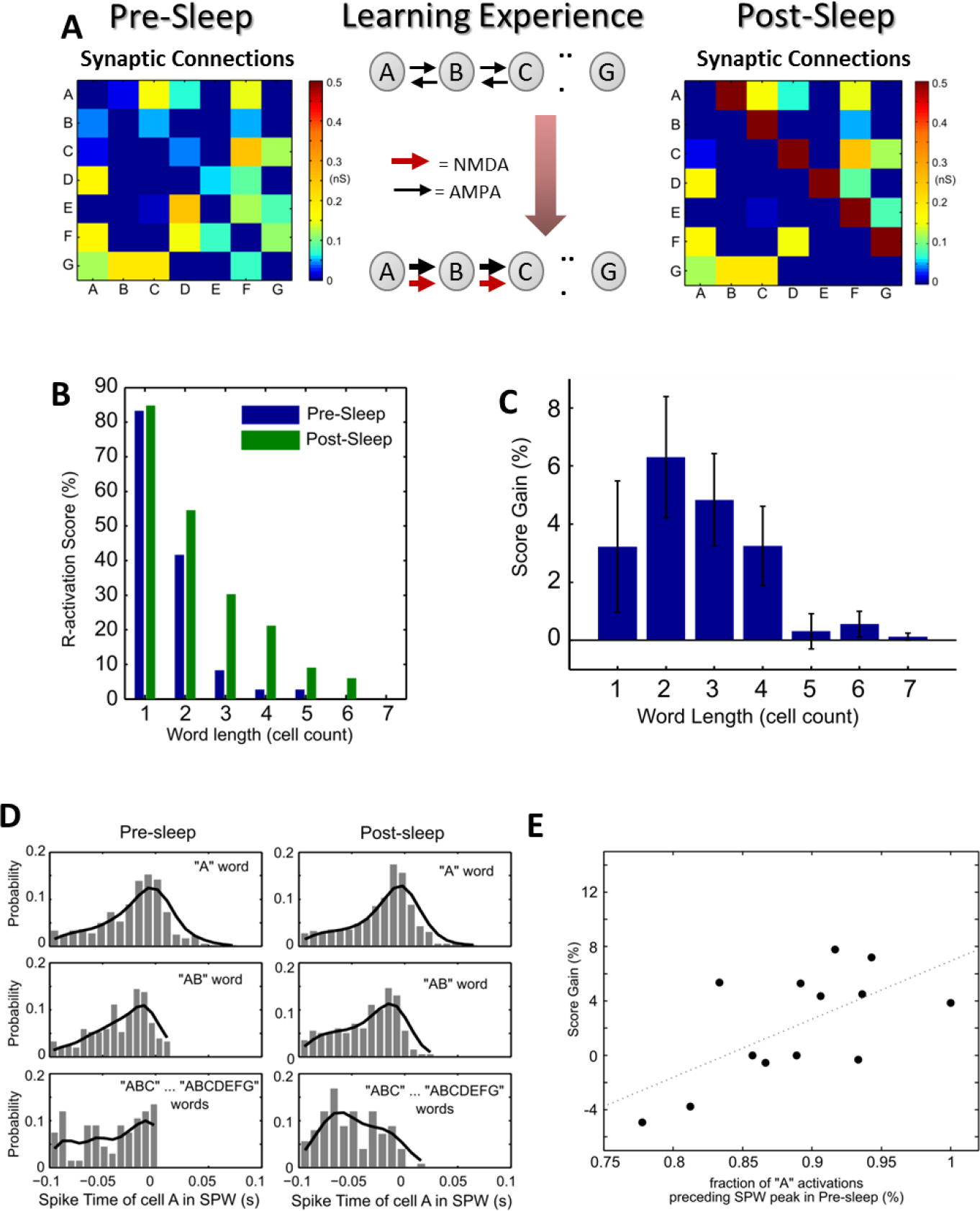
Learning extends word length reactivation. **A.** Changes in synaptic connections introduced in the paradigm. Drawing in the middle shows the manipulation we introduce to synaptic connectivity. AMPA synapses favoring the correct reactivation order were strengthened, while NMDA synapses favoring the word order were introduced. AMPA synapses promoting spiking in order opposite to the word order were removed. Matrices show the spike time difference among cells which constitute the ABCDEFG word in one example simulation, both in Presleep and in Post-sleep. **B.** Learning increases word R-activation at many lengths in one example simulation. Plot shows the R-activation scores for words of increasing lengths (“A” =1, “AB” =2, “ABC”= 3, etc.) both in Pre-sleep (blue bars) and Post-sleep (green bars).**C.** across 14 simulations, the plot shows the differences between Post-sleep R-activation score and Pre-sleep R-activation score of words of increasing lengths, averaged. Error bars show standard error. **D.** Length of word reactivation depends on timing of word initiation during the SPW. For each word activating during ripples, the time of spike of the first cell “A” (with respect to the peak of the sharp wave) is found and plotted against the length of the word. Near each word length dots, the error bar identifies the mean and standard deviation of the all the dots corresponding to words of the same length. Note that the length of an activating word seems related to the earliness of its first spike during the sharp wave. **E.** Learning-induced gain in word length depends on timing of first cell in Pre-sleep. For each simulation, the fraction of ripple reactivations of cell “A” with timing 12ms or more before the SPW peak (x-coordinate) is plotted against the total score gain for words of length 4 and above (i.e. summing the values of panel C for lengths 4 to 7). The linear fit shows a positive correlation (linear correlation r=0.6327 with significance p=0.0203). Note that in this figure we have discarded the data point (0.9667, 30.81) deemed an outlier because of its extremely high Score Gain.

To quantify the effect of these synaptic changes on the R-activation of cells in the sequence, we considered progressively longer sub-words within the total word length, always starting from cell “A”. Hence, we looked at the word “A” (length 1), “AB” (length 2), “ABC” (length 3), “ABCD” (length 4), etc. all the way to the full ABCDEFG word (length 7). For each word considered, across the different lengths, we found the percentage of SWR events in which the whole word was reactivated in the correct order, and called that the R-activation score of each word. We could then compare the R-activation scores of words of length 1 to 7 in Pre-sleep and Post-sleep. In Fig 3B, we show one example of the outcome of one simulation (Pre-sleep and Post-sleep): as it is intuitively necessary, the R-activation scores are lower for longer words, as they include the shorter ones. In this case, our artificial synaptic manipulation resulted in larger R-activation during Post-sleep of word of all lengths but the full one, meaning the last cell never spiked at the end of the whole sequence. It is to note that this measure is strict in the sense that it does not count the reactivation of partial words in the sequence unless it represents a chunk from the beginning (i.e. reactivation of BCDE in a SWR is not counted). Using this measure we could subtract the Pre-sleep R-activation from the Post-sleep R-activation scores to obtain a Score Gain for the R-activation of words of increasing lengths. Across 14 simulations, we found that artificially manipulating a small number of synapses promoted the R-activation of the sequence at all word lengths during Post-sleep (Fig 3C).

When studying the R-activation of triplets of cells (Fig 2) we had found that inputs to the first cell in the triplet could influence the synaptic-induced Score Gain. For a longer sequence of spikes the role of the first cell in the sequence was likely to still be relevant. In particular, we reasoned that the timing of the first spike in the sequence would affect the possible word length (within the sequence) which could reactivate in a given SWR: if the first spike happened too late in the SWR, only a small portion of the sequence could reactivate within the event duration. In fact, when we considered all sub-words reactivations across all 7 possible lengths, we could see that the timing of the first spike in the sequence (the spike of cell “A”) was widely distributed on both sides of the SWR peak for words of length 1 or 2 (“A” and “AB” case in Fig 3D), implying that for very short words the first spike could be late or early in the SWR event. For words of length 3 and higher, we found that the timing of their first spike was strongly placed to the left of the SWR peak, meaning it had to be preceding it (Fig 3D). This property (the earliness of first spike in longer reactivations) was true for both Pre-sleep and Post-sleep SWR simulations (Fig 2D), and the distributions did not look different in Pre vs Post-sleep, meaning our plasticity manipulation did not result in a generalized anticipation of the first cell spike. Hence, whenever our plasticity manipulation induced a large Score Gain for long sub-words, it had to prolong the length of a reactivated short sub-word which started early enough within a SWR. Based on this idea, we compared for each simulation the total Score Gain for words of length 4 and above with the fraction of the total cell A spikes during Pre-sleep which happened early in a SWR. Fig 3E shows the scatter plot of such comparison, confirming a statistically significant positive correlation.

### Experience-Related Learning: from rat trajectory to new synaptic connections

The next step in our modeling effort was to introduce an explicit relationship between a spatial learning experience and changes in synaptic connections, so that to create a virtual machinery in which a “rat” alternates spatial experiences (which can trigger learning among the network synapses) and sleeping experiences (which will show spontaneous reactivation during SWR activity). While modeling efforts to relate trajectory exploration to specific spiking of cells in a biophysical model have been developed (Kropff and Treves 2008, Burak and Fiete 2009, Giocomo, Moser et al. 2011), the specific mechanisms by which spatial perception results in place cell activity are currently under investigation, and are not the main subject of our study. Hence, we chose to not design a network spiking model of the awake state, but instead to introduce a phenomenological model of place cell spiking activity as our awake/learning state.

To model the learning phase of the “virtual rat” we follow a paradigm used to study episodic memory processing (Jones, Bukoski et al. 2012, Jones, Pest et al. 2015). We started by defining a 2-dimensional virtual enclosure (20×20 cm in size) and distributing 81 identical ‘place fields’ along the surface: 2-D Gaussian bumps of integral 1 with peaks evenly spaced, as shown in Fig 4A. In this virtual exploration enclosure, we introduced 8 locations of interest, marked with blue diamonds above the place fields in Fig 4A. These locations could be of interest to the virtual rat because there could be feeders or other types of reward placed at any or all of these locations. In this virtual enclosure, we represented a learning task consistent with the experiments which drove this model (Jones, Bukoski et al. 2012, Jones, Pest et al. 2015): we assigned to 3 of the feeders a special relevance (circled in yellow in Fig 4A) and we made the virtual rat explore 3 of these feeders in a fixed order, defining a learning trajectory (yellow arrows in Fig 4A). A learning experience was then constructed by a string of locations to be reached in time, where the 3 ordered learning targets were interspersed with 3 randomly chosen (varying each time) locations among the remaining 5. This produced a space exploration experience on the virtual enclosure *(x_t_,y_t_)*. As the virtual animal traveled in the enclosure, it traversed a number of place fields, triggering spikes with probabilities drawn from a Poisson process with rate given by the place field spiking probability. This is consistent with experimental measures which have shown that place fields can be reconstructed from the locations of spike times of place cells, and can be fitted with 2-D exponentials (O'Neill, Senior et al. 2006). Hence, for each exploration experience *(x_t_,y_t_)*, we considered spikes from 81 virtual cells encoding the place fields assigned to them. One example of such trajectory-induced spiking activity is shown in Fig 4B, where the spike times of a subset of all place cells are shown and shaded in gray are the times in which the virtual animal was completing the assigned relevant trajectory across the 3 rewarding feeders (in yellow in Fig 4A). In red we marked the spikes of cells with place fields along the straight path connecting the 3 relevant feeders, which can be seen spiking in order during the shaded times.

**Figure 4.**
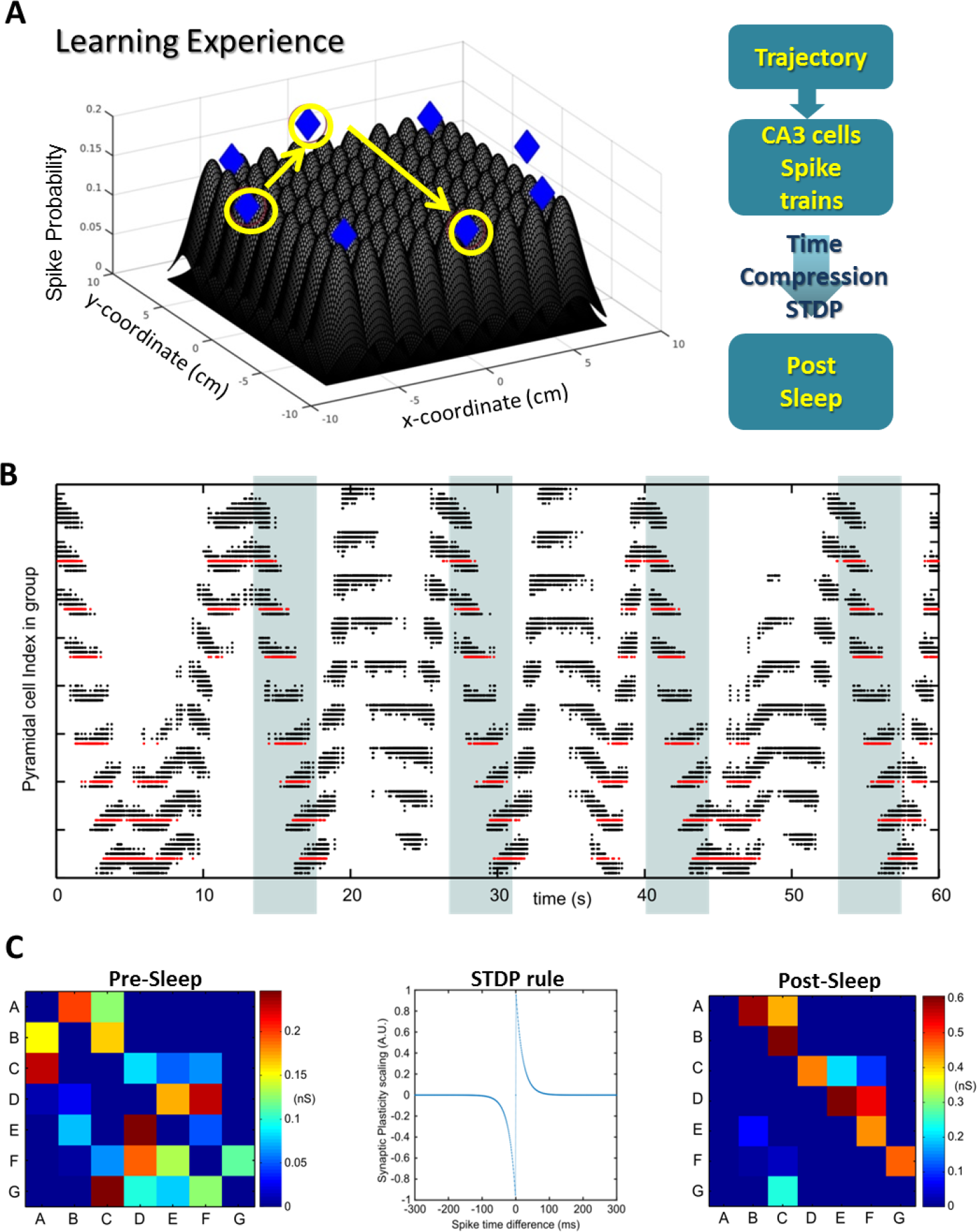
Experience-Related Learning: from rat trajectory to new synaptic connections. **A.** Representation of how a trajectory in space creates artificial spike times for selected CA3 cells. The available space is tiled by regions over which place fields are defined. Each field is assigned to a specific cell. The field defines the relationship between the rat coordinates in space and the probability of a cell to spike. The diagram on the right shows the procedure connecting a trajectory experience to a change in synaptic connections.**B.** Example of spiking of cells during learning experience. In the rastergram, each dot represents a spike time (x-axis) of a given cell (y-axis). In red are the spikes of cells subsequently used to identify a spiking sequence aBCDEFG for replay analysis (“trajectory cells”). In the shaded times, the “virtual rat” is exploring the sequence of locations to be learned (“trajectory”). In between times the animal is exploring randomly selected locations. **C.** Synapses between trajectory cells (red in B) before and after applying trajectory-driven plasticity (matrices on either side). Middle plot: representation of the function mediating the STDP rule used to connect spike times (as in B), after time-compression, to synaptic changes.

In an actual experiment, place cell spiking during exploration would happen within low frequency (3-8 Hz) theta oscillations in the LFP (Buzsaki, Buhl et al. 2003), and would be interspersed with awake SWR events, within which the recently learned sequences are known to reactivate, in a time compressed manner (Atherton, Dupret et al. 2015, Buzsaki 2015). It is hypothesized that synaptic plasticity within cells spiking during the awake SWR events is mediated by spike-time dependent plasticity (STDP) mechanisms, which strengthen synapses according to the relative spike timing of the pre-synaptic and post-synaptic cell. Specifically, if the pre-synaptic spike precedes the post-synaptic spike of a short time the connection is maximally strengthened, while if the time gap is wider the strengthening is a lot smaller. Vice-versa, when a post-synaptic spike precedes a pre-synaptic spike the connection is weakened. Hence, to model the change in synaptic connection strengths induced by a given learning/exploration experience we time-compressed the awake spike times by a factor of 10 and used a classic STDP rule to quantify the synaptic changes occurring between cells due to the spike times in the learning phase (see the middle panel in Fig 4C, and Computational Methods).

This portion of the setup completed the connection of an exploratory trajectory to spike times of a group of cells, and then to the computation of the STDP-induced rescaling of synaptic connections between those cells. This awake virtual structure had to be connected with our biophysical model of sleep SWR activity for the three-stage paradigm of Pre-sleep, learning and Post-sleep to be completed. By assigning each place field within the virtual enclosure to a given cell in the CA3 network, we could bridge the gap between awake virtual model and biophysical model of sleep. For each place cell in the enclosure, we selected one cell in our CA3 network, which was randomly assigned within a range of cell indexes. The range of CA3 pyramidal cells was selected to have a high likelihood of co-activation during SWR; since SWR are localized within the network topology, the 81 indexes were assigned to a range of cells of 100 indexes (details in Computational Methods). Since the cells belonged to the CA3 network, a specific synaptic strength value was already known from the Pre-sleep SWR activity simulation (see in the left panel of Fig 4C one example). The effect of STDP on virtual synaptic connections which resulted from the awake computation was then assigned offline to the synaptic connection strengths between the corresponding CA3 cells, hence completing the flow diagram shown in Fig 4A. One example of the resulting difference in synaptic connections between Pre-sleep and Post-sleep that was induced by this procedure is given in Fig 4C. Note that for each simulation we identified a sub-group of 7 cells among the 81 CA3 cells selected, to be the ones that host the spike sequence which is ‘recorded’ during our virtual experiment, and hence whose change in R-activation between Pre-sleep and Post-sleep was the subject of our analysis.

### Experience learning increases trajectory reactivation in Post-sleep

Once we had this explicit model of the Pre-sleep, learning, Post-sleep paradigm, we could observe the different R-activation of cells with place fields along the target trajectory in the two sleep epochs. In *in vivo* experiments, it is known that awake learning promotes reactivation in the subsequent sleep, so we first tested if our model satisfied this requirement. Since once again it was a list of 7 cells, we used the same measure of R-activation developed in the simplified learning case (when we only manually changed some synapses, Fig 3). Fig 5A shows one example of the R-activation of the 7 “trajectory” cells in Presleep and Post-sleep, for each word of increasing length, all starting from the first cell (“A”, “AB”, “ABC”, etc.). For each word length, we computed its Score Gain as the difference between Post-sleep and Pre-sleep R-activation scores and Fig 5B shows that across 14 simulations we had an increase in R-activation at many lengths, although the longest sub-words proved difficult to obtain.

**Figure 5.**
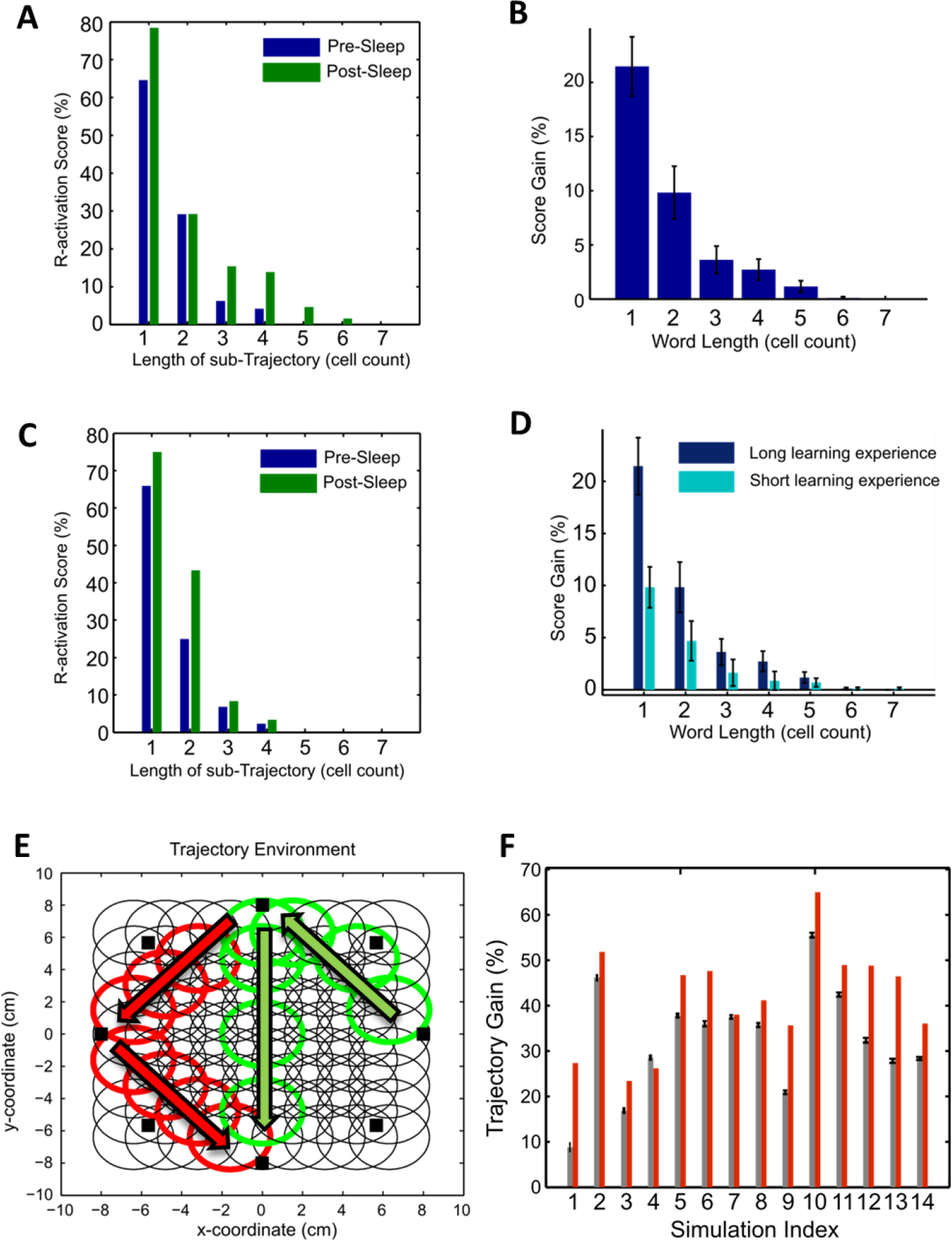
Experience learning increases trajectory reactivation in Postsleep. **A.** Trajectory learning increases sequence reactivation during sleep. The R-activation score of sequences of trajectory cells of increasing lengths (e.g., 1=“A”, 4=“ABCD”) in one example of Pre and Post-sleep simulations. **B.** Score Gain (Post-sleep score minus Pre-sleep score) for the cell sequences of increasing lengths. Bar shows mean value across 14 simulations, error-bars mark the standard error. **C.** Shorter learning experience results in reduced gain of sequence R-activation. Example of R-activation score of sequences in a Pre and Post-sleep simulation with trajectory learning time reduced to one half. **D.** Score Gain across sequence length in the case of long vs short learning experience. Note that at every length the increase in R-activation Post-sleep compared to pre-sleep is higher for stronger learning. Error-bars mark the standard error of the mean across 14 simulations. **E.** Example of the circular enclosure and two possible spatial trajectories: the trajectory which is learned in the task (in red) and one that is not learned in the task (in green). To compare the efficacy of learning on Score Gain we tested 336 possible trajectories (green) for each simulation against the red trajectory. **F.** For each simulation (n=14) we compare the Pre-sleep and Post-sleep R-activation of different trajectories. The red bar shows the Trajectory Score Gain for the learned trajectory, and the gray bar shows mean and standard error of the Trajectory Score Gain for the other 336 trajectories tested, which were not learned but were possibly visited in the task epoch.

In the virtual rat setup, we proceeded to study the role of different learning experiences on the R-activation changes between Pre-sleep and Post-sleep. For each simulation in Fig 5B, we had a spatial experience trajectory (performed during the learning phase). We repeated each simulation experiment using only half of the spatial experience trajectory. Since the exploration of the regions of interest leading to sequence learning was uniformly distributed along the spatial experience, this paradigm can be thought of as roughly halving the amount of training that the virtual rat received on the relevant feeders. We labeled the two training conditions “long learning experience” and “short learning experience”. Even with reduced learning, we still expected the plasticity induced by the spatial exploration experience to promote the R-activation of the cell sequence in Post-sleep. In Fig 5C we show one example of Post-sleep and Presleep R-activation scores in the short learning experience, confirming this expectation. However, reducing the learning time affected the Post-sleep R-activation: across all our simulations, the Score Gain of the short learning experience was smaller than the Gain for the long training (Fig 5D).

In our learning experience the virtual rat visited a sequence of 3 specific targets followed by 3 random targets, and then again the 3 specific targets, followed by 3 new random ones, etc. Hence, the path covered by the spatial learning experience involved in general spiking from any or all of the 81 CA3 cells with place fields tiling the spatial enclosure. We decided to compare the R-activation of a number of other possible trajectories, connecting any 3 feeders/targets in the enclosure, to the R-activation of the “learned trajectory”. In Fig 5E we show a view of the virtual enclosure from above, with circles representing the half-height of the place fields of the 81 pyramidal cells assigned to a given simulation. In red, we emphasize the path connecting the learned trajectory and the place fields of the cells which compose the spiking sequence “ABCDEFG” used in all quantifiers of R-activation scores above (Fig 5 A-D). In green, we mark one example of a different trajectory which was not the learning trajectory and of place fields of cells along its path. We collected a 7-cell long sequence for each possible 3-target trajectory given by the 8 target enclosure (a total of 336 samples), and for each sequence we found the Pre-sleep and Post-sleep R-activation for sub-words of increasing lengths. Since we wanted to compare learned and non-learned trajectories, we introduced a single number Trajectory R-activation Score as follows. For each length of a sub-word (from 1 to 7) we considered representative of a reactivation of a sub-word of length n all the ordered strings of letters of length n showing immediate neighbors within the main (7-letters) word. For example, for a sub-word of length 4, the representatives would be “ABCD”, “BCDE”, “CDEF” and “DEFG” (but “ACEF” would not qualify). The percentage of SWR in which any of the representatives of a sub-word of length 4 was spiking gave the R-activation score for length 4. A Trajectory Score in a given simulation (Pre-sleep or Post-sleep) would then be defined as the sum of all R-activation scores across all length (1 to 7). The difference between the Post-sleep Trajectory Score and the Pre-sleep Trajectory score then produced a Trajectory Gain. When the average Trajectory Score across all trajectories within the enclosure was compared with the learned trajectory, the learned trajectory scored above average in all but one simulation. Hence, our behavior driven synaptic plasticity paradigm could enhance specific trajectory reactivation with high reliability. This type of test would have been impossible in the simplified learning setup where we artificially changed only very few synapses in the network but no cell spiking activity was explicitly considered as coding for something specific in the awake epoch.

## Conclusion

In this work, we used computer models to study the impact of changing synaptic connectivity within the hippocampal CA1-CA3 network on the cell reactivation properties. We first introduced limited targeted synaptic changes between selected CA3 pyramidal cells, representing the effects of a learning experience in between two successive simulations of SWR activity, building a sequence of Pre-sleep, Learning and Post-sleep. We then expanded the learning epoch to represent explicitly a spatial exploration and path learning task, and applied STDP to trajectory-driven spiking to build a relationship between learning and connectivity, and hence study how learning can affect reactivation during sleep. For the sleep epochs, we used our previously developed model of SWR activity in CA3 and CA1 hippocampal regions. In this new work, we studied reactivation in the CA3 network component.

Specifically, for a given triplet of cells (Fig 1 and 2) representing part of the experience that was learned, we manually strengthened excitatory (AMPA) synapses favoring the triplet R-activation order, removed synapses opposing it, and introduced some NMDA connections (again favoring the triplet order) (Fig 1). Such manipulation in general resulted in a higher R-activation of the ABC triplet in the Post-sleep compared to the Pre-sleep (Fig 2). We found that the R-activation gained in the Post-sleep compared to Pre-sleep depended on the Pre-sleep score: triplets with lower Pre-sleep scores reached higher gains. Furthermore, our study highlighted a role for input (synaptic or increased intrinsic excitability) to the “first” cell of the sequence in promoting Post-sleep sequence reactivation, and in particular how timing of the first cell is crucial to the length of a sequence to be reactivated during a SWR (Fig 3).

The design of a full “virtual rat learning experience” model was inspired by rat behavioral paradigms used to study hippocampal ripple replay in an open enclosure (Fig 4). We designed a trajectory in space with a high amount of repetitions for three specific locations interspersed with varying groups of three random locations. We selected a group of CA3 pyramidal cells (81) and assigned to each of them a place field: a region in space where they were more likely to spike. Hence, for a given trajectory we could use Poisson processes to assign to each of the 81 cells spike times along the trajectory. We used these spike trains to calculate the effects of spike-time dependent plasticity (STDP) along the trajectory, and changed AMPA and NMDA synaptic connections in the whole CA3 network according to the resulting STDP (Fig 4). This method was again used to represent a learning experience in between two sleep simulations, and 7 cells with place fields along the trajectory were selected to represent the memory of the trajectory. When comparing R-activation in Pre-sleep and Post-sleep (Fig 5), we found that synaptic plasticity driven by the learning experience increased the spontaneous R-activation of the memory in CA3 during ripples, that such gain was dependent on the length of the learning experience and that while all cells involved in the learning process received changes in their synaptic connectivity the sequence representing the learned trajectory showed a larger gain than an exhaustive sample of other possible trajectories in the same enclosure.

Combining what we learned from the simplified synaptic change conditions with what we learned from our virtual rat learning experience, our study suggests the following hypothesis: that within the hippocampal system the choice of pyramidal cells that are recruited to encode for a specific task actively learned (a proxy for a human declarative memory) cannot be completely randomized. Instead, the pool of cells which are ready to be used for learning is tightly connected to the specifics of their reactivation activity in the previous night of sleep. In other words, the learning process operates in interaction with the network substrate, not independently of it. Thus, the synaptic changes mediating the selection of specific pyramidal cells for ordered co-reactivation during a Post-sleep event can be minimal, if the choice of cells used to code for the learning process is efficient. Specifically, the model predicts that a minimal number of synaptic adjustments can promote the largest increase in R-activation during Post-sleep for cell assemblies with a relatively low Pre-sleep R-activation score and with the first cell in the group showing early spiking during SWR in the Pre-sleep epoch.

We consider the model we present here a prototype which can be expanded to analyze and shape hypotheses related to a number of open questions, for example: 1) the influence of hippocampal reactivation on cortical reactivation can be studied in this model when connecting it to thalamo-cortical models of sleep rhythmic activity (Bazhenov, Timofeev et al. 2002, Krishnan, Chauvette et al. 2016), together with the role of cortical input in hippocampal SWR replay; 2) which synaptic changes are performed in the hippocampus when learning competing (interfering) memories, and how they can affect sleep-dependent consolidation of interfering memories (Mednick, Cai et al. 2011, Wei, Krishnan et al. 2016, Wei, Krishnan et al. 2016) 3) the model can be expanded to introduce plasticity to interneurons, and physiological awake spiking activity in the whole CA3-CA1 network (characterized by theta-gamma rhythmic interaction during active exploration (Kopell, Börgers et al. 2010)) to study the relationship between the theta phase spiking of a sequence and its reactivation during sleep SWR (Mizuseki, Royer et al. 2012, Grosmark and Buzsaki 2016).

This study suggests an explicit mechanism for STDP-mediated learning to interact progressively with SWR across one sleep-learning-sleep cycle, and can be interpreted within the broader problem of learning progressively new things across multiple learning/consolidation (awake/sleep) cycles. One hypothesis is that sleep dependent memory consolidation happens when reactivation of cell sequences in the hippocampus enables cortical reactivation, which in turn promotes plasticity within cortex, ultimately transferring the encoding of the memory information from the short-term storage of the hippocampus to the long-term distributed storage of cortex. It is known that across many days retrieval of a specific memory becomes progressively independent from hippocampal reactivation (Rasch and Born 2013). Our work suggests that as hippocampal sequences become less necessary to the memory consolidation process of a specific sleep cycle, they will reactivate less in SWR and therefore will become better suited to be used for coding new experiences in the following day. This evokes a scenario of cell assemblies “ranked” by their co-reactivation during Pre-sleep, where some will be used for learning, and hence increase their SWR co-activation in Post-sleep while the ones not used for learning will reactivate even less in Postsleep. This progressive ‘degradation’ of a memory representation in the hippocampus during sleep is generating a pool of readily available cell assemblies to use for coding the next day.

## Computational Methods

### Sleep epochs network model

#### Rationale

We started with our previously developed (Malerba, Krishnan et al. 2016) network of CA1 pyramidal and basket cells and constructed a network of pyramidal and basket cells to represent CA3 activity, then built Schaffer Collaterals projections from CA3 pyramidal cells to CA1 pyramidal cells and interneurons. We used equations of the adaptive exponential integrate and fire formalism (Brette and Gerstner 2005, Touboul and Brette 2008), which can show bursting activity (like CA3 and CA1 pyramidal cells (Andersen, Morris et al. 2006)) or fast-spiking activity (like basket cells (Andersen, Morris et al. 2006)) depending on their parameters (Brette and Gerstner 2005). CA3 pyramidal cells are allowed a stronger tendency to burst in response to a current step input by having a less strong spike frequency adaptation than CA1 neurons, consistently with data (Andersen, Morris et al. 2006). For simplicity, all cells belonging to the same population had the same parameters (specified in the following section). To introduce heterogeneity, every cell receives a different direct current term (selected from a normal distribution), together with an independent Ornstein–Uhlenbeck process (OU process) (Uhlenbeck and Ornstein 1930), which can be thought of as a single-pole filtered white noise, with cutoff at 100Hz. This noisy input is added to take into account the background activity of the cells which we did not explicitly model in the network. The standard deviation of the OU process controls the size of the standard deviation in sub-threshold fluctuations of cell voltages, and is a parameter kept fixed within any cell type. Once the parameter tuning is in effect, the cells (even when disconnected from the network) show fast and noisy sub-threshold voltage activity, and their spikes are non-rhythmic, driven by fluctuations in the noise input they received, which is called a noise-driven spiking regime, rather than a deterministic spiking regime, and is representative of *in vivo* conditions (Fernandez, Broicher et al. 2011, Roxin, Brunel et al. 2011, Broicher, Malerba et al. 2012).

Cells are arranged within a one-dimensional network in CA3, and connectivity within CA3 is characterized by each cell reaching other cells within a third of the network around them, which is consistent with anatomical estimates (Li, Somogyi et al. 1994). For pyramidal to pyramidal cells connections, the probability of synaptic contact within this radius of one third was higher for neurons closer to the pre-synaptic cell and decayed for neurons further away. Details of all network connections are introduced in the Connectivity section. Intuitively, the highly recurrent connections between pyramidal cells in CA3 has a gradient in density that induces a convergence/divergence connectivity fairly uniform across all CA3 pyramidal cells, which represents the homogeneity of CA3 pyramidal cells arborization within the region. This connectivity represents the highly recurrent pyramidal connections in CA3 without introducing special hubs of increased excitatory recurrence in any specific location in the network.

#### Equations and parameters

We model SWR activity in the hippocampus using a network of 240 basket cells and 1200 pyramidal cells in CA3, 160 basket cells and 800 pyramidal cells in CA1. The ratio of excitatory to inhibitory neurons is known to be approximately 4 (Andersen, Morris et al. 2006) and since in our model we did not introduce any of the numerous hippocampal interneuron types but for basket cells, we apply that ratio to the pyramidal to basket cell network. This ratio also favored the ability of the network to support a background disorganized spiking regime, where excitatory and inhibitory currents were able to balance each other (Atallah and Scanziani 2009). For each neuron, the equations are

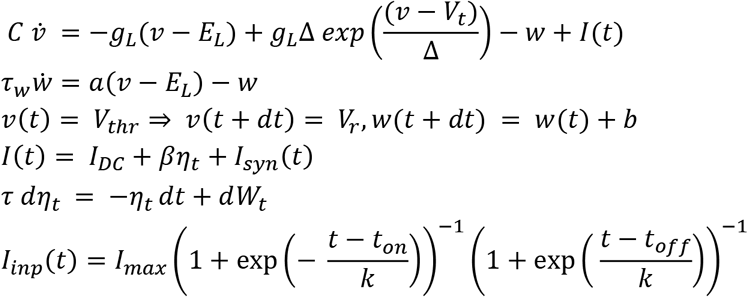

CA1 cells parameters are reported in (Malerba, Krishnan et al. 2016), and CA3 cells parameters were as follows. Pyramidal cells parameters: C (pF) = 200; g_L_ (nS) = 10; E_L_ (mV) = −58; A = 2; b(pA) = 40; Δ (mV) = 2; τ_W_ (ms)=120; V_t_ (mV) = −50; V_r_ (mV) = −46; V_thr_ (mV) = 0. Interneurons parameters: C(pF) = 200; g_L_ (nS) = 10; E_L_(mV)= −70; A = 2;b(pA) = 10; Δ (mV) = 2; τ_w_(ms) =30; v_t_(mV) = −50; V_r_(mV) = −58; V_thr_(mV) = 0.

The coefficients establishing noise size were β = 80 for pyramidal cells, β =90 for interneurons. DC inputs were selected from Gaussian distributions with mean 24 (pA) and standard deviation 30% of the mean for pyramidal cells in CA3, mean 130 (pA) and standard deviation 30% of the mean for CA3 interneurons, mean 40 (pA) and standard deviation 10% of the mean for CA1 pyramidal cells and mean 180 (pA) and standard deviation 10% of the mean for CA1 interneurons.

Synaptic currents were modeled with double exponential functions, for every cell *n* we had *Isyn* (*t*)= 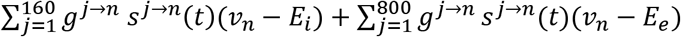, where *E_i_* = −80 mV and *E*_*e*_ = 0 mV, and 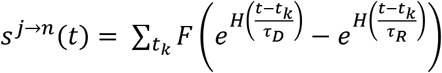, where *t*_*k*_ are all the spikes of pre-synaptic cell *j*.

In this equation, *F* is a normalization coefficient, set so that every spike in the double exponential within parentheses peaks at one, and *H*(·) is the Heaviside function, ensuring that the effect of each pre-synaptic spike affects the post-synaptic current only after the spike has happened. The time scales of rise and decay (in ms) used in the model were as follows (Bartos, Vida et al. 2002, Cutsuridis, Graham et al. 2010, Taxidis, Coombes et al. 2012, Malerba, Krishnan et al. 2016). For AMPA connections from pyramidal cells to pyramidal cells: τ_R_=0.5, τ_D_=3.5. For AMPA connecions from pyramidal cells to interneurons: τ_R_=0.5, τ_D_=3. For GABA_A_ connections from interneurons to interneurons: τ_R_=0.3, τ_D_=2. For GABA_A_ connections from interneurons to pyramidal cells: τ_R_=0.3, τ_D_=3.5. For NMDA synapses between CA3 pyramidal cells, which we add in the learning paradigm and are hence present in Post-sleep network activity simulations, τ_R_=9, τ_D_=250 (Andersen, Morris et al. 2006).

#### Connectivity

The CA3 network was organized as a one-dimensional network. For connections from a CA3 pyramidal cell to the other CA3 pyramidal cells, we first considered a radius (of about one third of the network) around the presynaptic cell, and the probability of connection from the presynaptic cell to any cell within such radius was higher for cells with indexes nearby the presynaptic cell and reduced progressively with cell index distance (Li, Somogyi et al. 1994). Specifically, we used a cosine function to shape the probability within the radius, and parameterized how fast with index distance the probability had to decay by using a monotonic scaling of the cosine phase: if x was the index distance within the network, y = arctan(kx)/arctan(k) imposed the decay probability p(y) = Pcos(4y), where P was the peak probability and k = 2 was a parameter controlling the decay of connection probability with distance within the radius. An analogous structure underlid the probability of CA3 pyramidal cells to connect to inhibitory interneuron in CA3 and for Schaffer Collaterals to connect a CA3 pyramidal cell to CA1 pyramidal cells (Li, Somogyi et al. 1994). To balance the relationship between feed-forward excitation from pyramidal cells to interneurons and feedback inhibition from interneurons to pyramidal cells, probability of connection from a presynaptic basket cell to a cell within a radius (about 1/3 of the network size) was constant at 0.7, for GABA_A_ connections to both CA3 pyramidal cells and interneurons. Within CA1 connectivity was all-to-all, with the caveat that synaptic weights which were sampled at or below zero caused a removal of a given synapse. As a result, most synapses between CA1 pyramidal cells were absent, consistently with experimental findings (Deuchars and Thomson 1996). To introduce heterogeneity among synaptic connections, synaptic weights for all synapse types were sampled from Gaussian distributions with variance (σ) given by a percent of the mean (μ). Parameters used in the simulations were (we use the notation Py3 and Py1 to denote pyramidal cells in CA3 and CA1, respectively and analogously Int3/Int1 for interneurons). Py3->Py3: μ = 34, σ = 40%μ; Int3->Int3: μ = 54, σ = 40%μ; Py3->Int3: μ = 77, σ = 40%μ; Int3->Py3: μ = 55, σ = 40%μ; Py3->Py1: μ = 34, σ = 10%μ; Py3->Int1: μ = 320, σ = 10%μ; Int1->Int1: μ = 3.75, σ = 1%μ; Py1->Int1: μ = 6.7, σ = 1%μ; Int1->Py1: μ = 8.3, σ = 1%μ; Py1->Py1: μ = 0.67, σ = 1%μ. It is to note that the mean (μ) declared was normalized by the total number of cells before the variance to the mean was introduced in the distribution. Since the CA3 and CA1 networks are of different sizes, a direct comparison of the parameter values or their magnitude across regions would not account for the effective values used in the simulations.

### Learning epochs models

#### Introducing targeted synaptic changes

##### “ABC” triplets analysis

In the CA3 network, we looked for pyramidal cells that had a chance to reactivate in a large fraction of spontaneous SWR. In our model, the probability of SPW participation among CA3 cells depends on network topology, and in particular looks bi-modal with peaks about cell index 350 and 750 (data not shown). Hence, we started from an interval of accepted cell indexes between 700 and 800. Three cells within this range were then chosen uniformly (using function randperm in Matlab, The Mathworks). For each Pre-sleep simulation (we had 15 samples, each 50s long), 20 sets of 3 cells were first randomly chosen and then cells within the sets were permuted, generating a set of 120 ordered triplets. From this pool, three triplets were chosen with the criteria that no two triplets could be a permutation of each other and that their R-activation scores in Pre-sleep were spanning a range of available scores (meaning one triplet was chosen with a low score, one with a medium score and one with a high score). Hence we had a pool of 45 triplets with Pre-sleep R-activation Scores roughly spanning the available range.

For each triplet of cells “ABC”, we manually modified some AMPA and NMDA synapses between the cells composing the triplet. The synaptic weights of AMPA synapses favoring the order of the triplet (A->B and B->C) were increased to the maximum AMPA synaptic strength value within the specific Presleep CA3 network (remember for every simulation the synaptic connectivity matrices were generated anew). Along the maximized AMPA synapses, we introduced NMDA synaptic connections with a fixed value, identical for all simulations (1.25nS, normalized by the total number of added NMDA synapses), which was found to be efficient at promoting co-activation during SWRs without enhancing the spiking activity of cells beyond physiological values. Finally, the weights of AMPA synapses which were favoring the reverse order of the triplet (C->B and B->A) were replaced by zeros. The new Post-sleep simulation was then run for 50s.

The R-activation Score was found using the spike times of the cells in the triplet. The spike times of the three cells were required to be between the start and end of the SPW event in CA3. The first spike of each cell was used to identify the order in which the cells spiked relative to each other. The R-activation Score of a triplet in a simulation was the percent of the total SWR in which the ordered triplet spiked (meaning each cell of the triplet spiked at least once and in the correct order). The Score Gain was found by taking the difference between the R-activation Score in Post-sleep and Pre-sleep. To verify whether the cells were spiking in the correct order, we also quantified the average spike time difference (STD) across triplets.

We analyzed the input received by the triplet “ABC” by using synaptic and intrinsic excitability modifications. For the synaptic modifications, an additional cell “D” was randomly chosen from the cell index range 700 to 800. This cell was added to the end of each of the ordered triplets from the 15 simulations. The same synaptic modifications in the NMDA and AMPA synapses were performed between the additional cell and the last cell of the triplet. The Score Gain between the triplet “ABC” and “BCD” was then compared to analyze the effect of altering the synapses between “A” and “B” on the replay of “BCD”. To confirm whether the input of the first cell influences the replay of the triplet, the intrinsic excitability of the first cell in the triplet was modified. The DC input of the first cell (*I_DC_* in the membrane voltage equation) was increased by 2pA (a small value, chosen to maintain the cells in a noise-driven spiking regime without driving them to an oscillatory bursting spiking regime). This intrinsic excitability was then doubled and tripled.

##### “ABCDEFG” sequences

We tested the effects of targeted synaptic modifications for longer cell sequences (7 cells, represented with the word “ABCDEFG”). To select 7 cells from our range of cell indexes between 700 and 800 in CA3, we proceed starting from a triplet and progressively adding one more cell at a time, as follows. The starting triplet was randomly chosen, with two requirements: 1) in Pre-sleep synapses between A->B and B->C were non-zero and 2) in Pre-sleep the R-activation Score of the triplet was between 4 and 10%. The next cell (“D”) was randomly chosen among the remaining cells in the 700-800 range with the same requirements as above applied to the new sub-triplet “BCD” (i.e. a synapse C->D was present in Pre-sleep and the BCD R-activation score was in the 4-10% range). The procedure of adding cells at the end of the list was iterated until a 7-letter list of cell indexes was populated. For these tests we used 14 simulations, 50s long. Since we did not analyze the CA1 cells, we only ran simulations of the CA3 sub-network in this part of the study.

To quantify the R-activation of sub-words in the sequence, the time of the first spike of any cell within each SWR was used. It is important to note that we effectively composed a list of 8 cells and then used the last seven of them as the ABCDEFG of our sequence, to possibly take into account the role of input to the first cell which we learned about in the triplet analysis. Furthermore, for cells which were later in a sequence (from cell D onward), we included their first spike time even if it happened up to 0.3s after the end of a SPW, to let the possible “tales” of a reactivating sequence be included in our analysis.

##### Modeling a virtual rat’s spatial learning experience

The “learning experience” comprised of a rat running across an artificial enclosure, which we tiled as a grid of place fields. The size of the enclosure was 16 cm by 16 cm and the grid was spaced with 0.1 cm gaps. Each place field was numbered from 1 to 81. Each place field represented a Poisson distribution of the cell’s firing rate as function of location, as a normal probability density function with center at one of the grid’s 81 locations and radius (standard deviation) 3cm. This radius was chosen to establish a sufficient overlap between place fields for the spiking activity to lead to significant STDP-mediated synaptic rescaling.

We chose a simple trajectory for the rat to learn. The rat had to move through 3 locations in the same order. These 3 locations, as well as the direction of the trajectory, were the same for all simulations. The rat then moved to 3 random locations that were not part of the learned trajectory. This counted as one repetition for the learning task. The rat moved between the locations in a straight line that was completed in about 2 seconds. Throughout the movement, each place cell spiked at a hundredth of a second.

When assigning a place field (1 to 81) to a CA3 cell within the sleep model network, we started from 7 place fields along the learned trajectory, and assigned to them the same 7 CA3 cells used for the simulations with targeted synaptic changes. The remaining 74 cells were randomly assigned within the 700-800 index range (excluding the cells which already received a field). Given a “learning experience” trajectory, we had 81 spike trains generated by their respective Poisson processes place fields, and used them to modify the synapses between the network cells which were assigned those place fields. We used Spike Time Dependent Plasticity (STDP) as the rule for strengthening or weakening AMPA and NMDA synapses. The spike times were compressed by factor of 10, to represent their reactivation during awake SWR leading to STDP-induced synaptic plasticity (Sadowski, Jones et al. 2016). Every synaptic connection strength g from cell A to cell B was then replaced by *g*+Δ*g* with 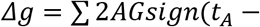. Where *g*_*max*_ = *A* ∗ *G* with A = 0.001 for AMPA synapses and 0.01 for NMDA synapses a scaling factor, G the maximum value of AMPA synapses in the given CA3 network, and G = 1.25nS for NMDA synapses. *t_A_* and *t_B_* are spike times for cells A and B, respectively.

Within a given “learning experience” the rat repeatedly visited the learning trajectory, followed by 3 random locations, which were newly re-selected with each iteration. We established the length of a given learning experience path to represent a long learning experience or a short one depending on the change induced in the synapses by the path. In fact, since the STDP effect is cumulative and every iteration results in incremental increase in the synapses favoring a target sequence ordered R-activation. The length of the experience path was determined by incrementally increasing the repetitions until the average strength of AMPA synapses between cells along the trajectory reached a threshold of 0.4nS. This value is close to the maximum synapse for all simulations used (the average maximum strength of the AMPA synapses over all 14 simulations was 0.517 with a standard deviation of 0.023). We chose a threshold value below the maximum AMPA synapse for each simulation because some synapses could significantly surpass the threshold and disrupt the spontaneous network activity.

## Acknowledgements

This work was supported by MURI grant (MURI: N000141612829 and N000141612415) to MB.

